# On the Impact of Smoking on The Oral-brain Axis: The Microbiome-Metabolism-Brain Interactions

**DOI:** 10.1101/2025.04.23.650308

**Authors:** Siti Maghfirotul Ulyah, Symeon Savvopoulos, Mohammad T. Albataineh, Herbert F. Jelinek, Haralampos Hatzikirou

**Affiliations:** Mathematics, Khalifa University, Al Sa’ada, Abu Dhabi, 127788, United Arab Emirates; Mathematics, Airlangga University, Mulyorejo, Surabaya, 10587, Indonesia; Department of Diseases’ Basic Sciences, College of Medicine, Yarmouk University, Shafiq Irshidat, Irbid, 21163, Jordan; Biotechnology Center, Khalifa University, Al Sa’ada, Abu Dhabi, 127788, United Arab Emirates; Department of Medical Sciences, Khalifa University, Al Sa’ada, Abu Dhabi, 127788, United Arab Emirates; Center for Information Services and High Performance Computing, Technische Univesität Dresden, Nöthnitzer Straße 46, Dresden, 01062, Germany

**Keywords:** Oral-brain axis, metabolism, microbiome, psychology, machine learning, dynamic system, mathematical model

## Abstract

Systemic interactions between distant organs, such as the oral-brain axis, remain largely unknown. This study aimed to shed light on a specific aspect of these oral-brain interactions, by focusing on the brain, in the form of psychological traits, the oral microbiome, and metabolism (BMM) with respect to the influence of smoking on the oral microbiota. We analyzed three datasets covering microbiome composition, psychological traits, and metabolic pathways in smokers and non-smokers. We used Canonical Correlation Analysis (CCA) to reduce the dimensionality of the three different datasets into scalar variables, and network analysis to identify the interactions between important features of the BMM system. Nonlinear regression revealed steady state interactions, showing tight correlations between metabolic pathways and microbiome composition. Smoking weakens the directed interaction between metabolism and psychological traits. Sensitivity analysis demonstrated the system’s robustness to small perturbations, but large disruptions could destabilize it, potentially leading to oral pathologies or psychological conditions. These findings highlight smoking’s complex role in BMM interactions and its implications for managing psychological conditions in smokers.

## 1 Introduction

Although anatomically separated, distant organs communicate with one another [1, 2]. For example, the gut-brain axis forms a network of connections across multiple biological systems, enabling bidirectional communication between gut bacteria and the brain [3]. Research on microbiome-Central Nervous System (CNS) interactions ranges from correlation studies to mechanistic explorations [4]. Both preclinical and human studies suggest that the gut microbiome plays a role in regulating depressive behavior [5, 6, 7, 8, 9], social behavior [10, 11, 12, 13], physical performance and motivation [14, 15, 16], and neurodegenerative diseases [17]. Another example of interconnection is the oral-gut-brain axis, which is characterized by anatomical communication, as the oral cavity and gastrointestinal tract continuously. The oral-brain axis is mainly regulated by the trigeminal nerve, whereas the gut-brain axis is primarily influenced by the vagus nerve [18].

An interaction in the oral-brain-axis was proposed in many studies. Research examining the link between experimental periodontitis and neurodegenerative disorders, such as Alzheimer’s disease (AD) and Parkinson’s disease, suggested that periodontitis in different models can disrupt the balance of the intestinal microbiota, compromise intestinal barrier integrity, activate immune responses in the gut, and contribute to neuroinflammation and neurodegeneration [19, 20, 21]. Malan et al. investigated the intricate connections between oral microorganisms, periodontal diseases, and mental health conditions. Their study revealed that both periodontal and mental health factors significantly shape the composition of the oral microbiome, with certain microbial taxa being linked to mental health, trauma, and overall well-being. In addition, the overall oral microbial composition was affected by several factors, with the greatest effect observed from smoking status [22]. Bowland et al. explored anthropological perspectives that integrate sociocultural, epidemiological, genetic, and environmental influences in shaping oral microbial ecosystems and their role in neuropsychiatric disorders (NPDs). Their findings indicated that oral microorganisms likely play a key role in the onset, progression, and manifestation of NPD symptoms through the oral-microbiota-brain axis [23].

Our study focused on investigating the interaction between psychological characteristics (for abbreviated as Brain), the oral Microbiome, and Metabolism (BMM). The oral microbiome, a crucial gateway with a distinct composition, is functionally connected to both the gut and respiratory microbiomes [24]. Additionally, the gut microbiota plays a significant role in behavioral disorders, among other health-related conditions [25].

Our objective is to understand the interplay between the oral microbiome, metabolic processes, and psychological characteristics, with a particular focus on smokers. We developed a tailored data-driven dynamical system model to account for the effect of smoking on the gut-brain axis. Recent research has explored the relationship between nicotine dependence and oral microorganism function in Middle Eastern populations, who are commonly known as heavy smokers [26]. Al Bataineh et al. identified significant compositional and functional changes in oral microbial communities, particularly among nicotine-dependent individuals, with associations to respiratory diseases and smoking cessation relapse [26]. Our study was based on these data and prospectively added psychological data to gain a better understanding of the effect of changing the oral microbiota on psychological health.

Previous studies have explored the relationship between microbes and the brain using various statistical and machine learning methods [27]. These include linear regression and analysis of variance (ANOVA) [28, 29, 30, 31, 32], amorphomic ANOVA [33, 34, 35, 36], partial correlation [37, 38, 39, 40], correlation-based triangular network analysis [41], linked independent component analysis (LICA) [42], spatial canonical correlation analysis (sCCA) [29], and parametric empirical Bayes (PEB) analysis [29]. Other studies have applied differential abundance testing methods, including sparse partial least squares discriminant analysis (sPLS-DA) [37, 36], linear discriminant analysis effect size (LefSE) [31], multivariate association of microorganisms with linear models (MaAsLin2) [31], and differential gene expression analysis based on a negative binomial distribution (DESeq2) [36]. Our approach moves beyond statistical and machine learning analyses by focusing on the potential dynamic behavior of the BMM system, providing a more comprehensive understanding of the underlying gut-brain interactions and their associations with psychological traits.

This paper is organized as follows: Section 1 provides the background and objectives of examining the effects of smoking on BMM interactions. Section 2 describes the data, variables, and methodology. Finally, Sections 3 and 4 present the results and discussion, respectively.

### Approach Rationale

In this study, the three datasets employed were the oral microbiome composition (Microbiota), metabolic pathway components (Metabolism) and psychological traits (Brain) for smokers and non-smokers data (Section 2.1). Our goal was to derive the interactions that drive the dynamics of the BMM network for smokers (s) and compare them with non-smokers (ns) (Figure 1). To identify differences between (s) and (ns), summary statistics were provided. Subsequently, Canonical Correlation Analysis (CCA) was employed to reduce the dimensionality of the three datasets into scalar coarse-grained variables (Section 2.2). This step allows us to map each dataset into a single scalar variable (covariate), which is a linear combination of the underlying features. The main idea behind this dimensionality reduction is to identify a system of equations that represent the BMM interactions. CCA also allows for identifying potential important features in each dataset. A network analysis and data clustering, provides further insights into the connections between important features of highly correlated covariate pairs (Section 2.3).

**Fig. 1:**
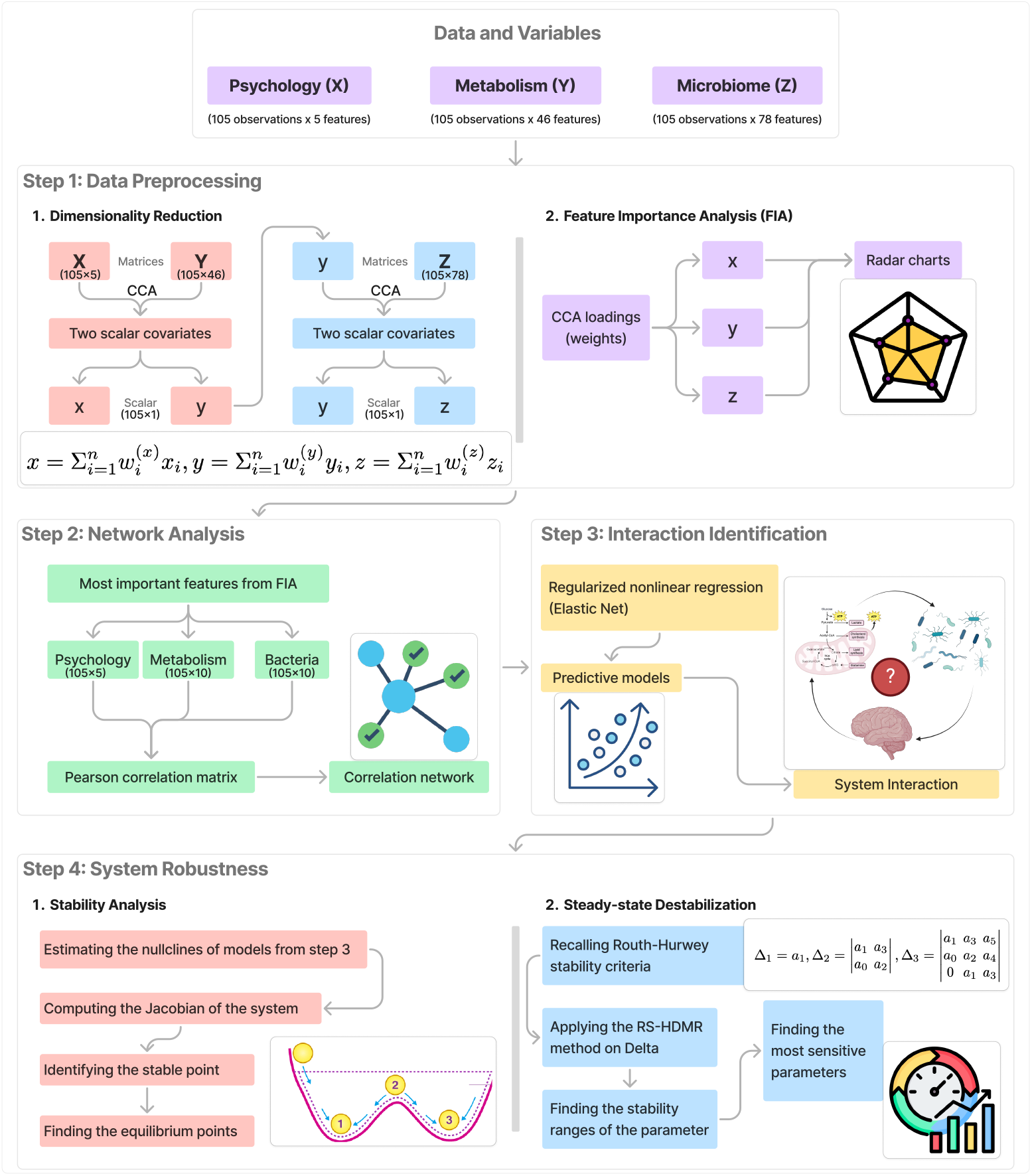
Schematic representation of our approach and the methodologies used, as explained in “Approach Rationale”. See Subsection 1 for further details.

Moreover, nonlinear regression was applied to derive relationships between the CCA-derived covariates (Section 2.4). These relationships can be viewed as steady state nullclines of the dynamics that dictate the Brain-Microbiota-Metabolism interactions. In examining how these interactions differ between non-smoker and smoker groups, it is expected that healthy physiological interactions operate close to equilibrium [43].

Finally, the steady states and the corresponding stability properties for smokers and non-smokers were calculated along with the parameter sensitivity. The latter provides insights into which parameters are important in perturbing the healthy steady state of non-smokers and potentially leading to various diseases (Section 2.5).

## 2 Materials and Method

This section describes the source and details of the data, data pre-processing and further analysis. These include dimensionality reduction and feature importance analysis, network analysis, predictive model and stability analysis. The summary of the method is described in Figure 1.

### 2.1 Data and Variables

A total of 105 samples, 55 smokers and 50 non-smokers, were drawn from individuals in the United Arab Emirates who agreed to participate in the research and signed a consent form after an introductory session explaining the research. There are three main variables, consisting of several indicators (features), as follows:

- **Brain:** Psychological traits 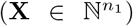 has *n*_1_ = 5 features (Extraversion, Agreeableness, Conscientiousness, Negative Emotionality, and Open Mindedness)
- **Metabolism:** Metabolic Pathways 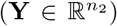 has *n*_2_ = 46 metabolic pathways features
- **Microbiota:** Bacteria Composition 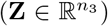 consists of *n*_3_ = 24, 925 species. A total of 99% relative abundances of the bacteria were taken and classified into 78 genera.

The psychological traits, including the five fundamental dimensions of personality, are often referred to as the five main personality traits [44, 45].

The detailed features of Metabolic Pathways and Bacteria Composition, consisting of bacterial genera, are in Tables A1-A2 (see the Appendix). These tables describe the notation of the 46 features of metabolic pathways and 78 genera in the composition of bacteria.

### 2.2 Data Preprocessing

Data preprocessing was the first step in identifying the unknown relation among variables in these data. In general, the pre-processing stage includes dimensionality reduction and feature importance analysis. The metabolic pathways and bacteria composition variables have a large dimension of features, while the sample size was only 105 observations. Due to the curse of dimensionality, it was difficult to draw any conclusions. Thus, CCA was employed. Before proceeding to CCA, the data were scaled from zero to unity.

CCA converts multivariate data into orthogonal variables known as canonical variates (CVs). Canonical variates represent linear combinations of the initial variables within each dataset, to maximize the correlation between the two datasets [46]. This method aims to reduce the dimension of each variable, which has several features, by considering the association among those three variables. The detailed steps are as follows:

1. Performing the first CCA between **X** (Brain) and **Y** (Metabolism), then extracting the first canonical variates, which are the linear combination of the features of each variable:

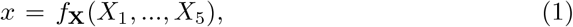

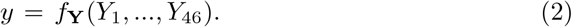
2. Performing the second CCA between **Z** (Microbiota) and the first canonical variates of **Y** (Metabolism), i.e. *y*, then extracting the first canonical variates of **Z** (Microbiota):

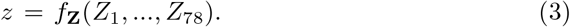 The three canonical variates are then further analyzed utilizing stability analysis (*x, y, z*).
3. Performing a feature importance analysis based on CCA loadings.
4. Testing the mean rank differences between smokers and non-smokers using the Kruskal-Wallis test on the original data of the ten most important features for each variable. Five percent significance level was applied here.

The two-stages CCA was conducted with some considerations: (i) the correlation between **Y** and **Z** (*ρ*_*Y Z*_) is very strong, but (ii) the correlation between **X** and **Y, Z** (*ρ*_*XY*_, *ρ*_*XZ*_) is rather low. Therefore, in the second stage (step 2), we run the CCA between **Z** and *y*, which accommodates a strong correlation with **X**. Using this setting, the covariates can capture the relationship among the three variables.

### 2.3 Network Analysis

A correlation network between the CCA important features was plotted to determine the linear association between the variables. This network was based on the results of the Pearson correlation test with 5% significance level that indicates the significant correlation among the most important features of the variables. The node size represents the number of relations in which a larger node means more association. Furthermore, we also identified the correlation between all the features and clustered the correlation into three groups, representing a weak to strong linear correlation between them. The *ward* hierarchical clustering method was applied here.

### 2.4 Interaction Identification

Nonlinear regression analysis was used to show how these variable interact. In turn, we assume that the expected value of the non-smoker population data corresponds to a stable, steady state of the system [43]. Thus, our regression system represents the steady state of the system’s potential dynamics. By applying linear stability analysis, we can identify the system’s stability properties.

Suppose that we have a system of ordinary differential equations as follows.

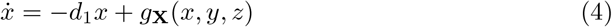

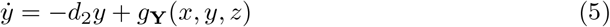

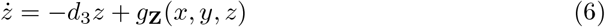

where *d*_*i*_ is the degradation coefficient for *i*=1,2,3. The nullclines of this system are:

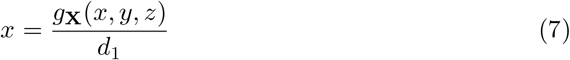

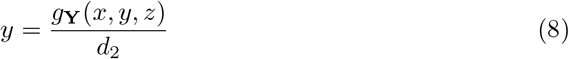

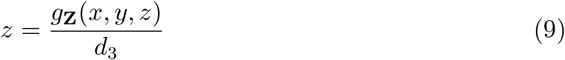

The right-hand side of Equations 7-9 will be denoted by **f**, where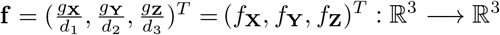.

Nullcline is a concept used for analyzing dynamical systems, especially for examining differential equations. It helps to visualize and understand the behavior of differential equation systems. Nullclines are the set of points in the phase plane where the first derivative of the variable with respect to time equals zero (e.g.,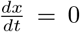). The intersections of the nullclines are equilibrium points [47]. These nullclines were estimated using regularized nonlinear regression as follows.

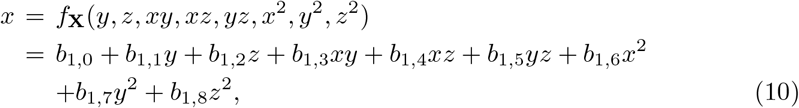

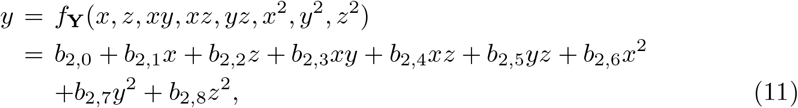

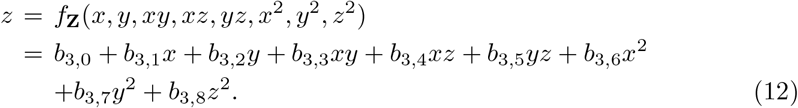

One should note that the covariates of the variables were used in this analysis. Thus, an elastic net algorithm must be applied as the regularization method because the covariates are highly correlated. Some parameters of the regularized nonlinear regression are *L*_1_*wt* = 0.3 and penalty = 0.02. *L*_1_*wt* is the L1 weight that ranges from zero to unity, where zero means pure Ridge regression and 1 represents LASSO.

### 2.5 System Robustness

For the stability analysis, the Jacobian of equation system (10 - 12) was calculated. Then, the stability of the point was assessed. After that, a global sensitivity analysis was performed using the random sampling - high dimensional mathematical representation (RS-HDMR) method to examine the sensitivity of the parameters in the system. RS-HDMR is essential to understanding the impact of complex model outputs on input factors. This method evaluates all input components simultaneously across their entire range of values, in contrast to local sensitivity analysis, which evaluates period changes in input parameters around a specific point. Models with non-linear input interactions are advantageously affected by this comprehensive approach. The primary advantage of RS-HDMR is its ability to allocate the model output uncertainty to the input factor uncertainty. This helps in identifying most critical inputs and necessitates meticulous measurement or administration. It also helps prioritize resources to understand and eliminate uncertainties in critical inputs [48]. RS-HDMR is indispensable in the fields of environmental modeling, engineering, and economics, where models exhibit intricate non-linear characteristics [48, 49, 50, 51]. RS-HDMR enhances the accuracy, reliability, and decision-making of models by quantifying the contributions of inputs to output variability. In general, local sensitivity analyses fail to account for input variable interactions; however, RS-HDMR can demonstrate the impact of changes in one element on another. Understanding and managing the uncertainty of the prediction of the model is essential for reliable and robust modeling, as demonstrated by RS-HDMR [48, 49, 50, 51].

RS-HDMR, which is analogous to the Sobol global sensitivity analysis technique, deconstructs the output of a model into components of increasing dimensionality [52].

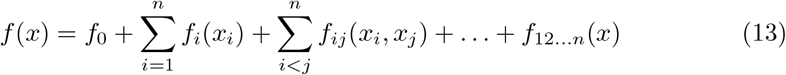

where:

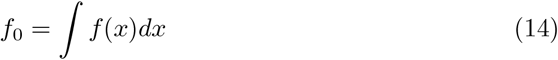

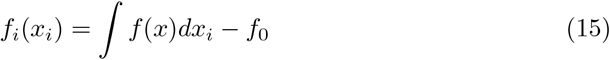

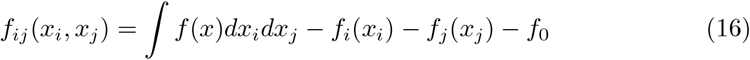

and *x* = [*x*_1_, *x*_2_, …, *x*_*n*_] is the input of parameters whose values change; *dx*_*i*_ is the product *dx*_1_*dx*_2_ … *dx*_*n*_; *dx*_*ij*_ is the same product without *dx*_*i*_ and *dx*_*j*_. The decomposition, known as ANOVA, generates distinct and independent terms and uncovers both primary and secondary effects to represent output using methods based on variance. The total variance *D* is [53]:

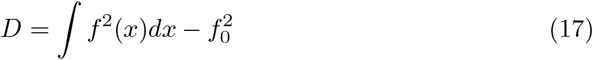

Further, the partial variances *D*_*ij*…_ are calculated by:

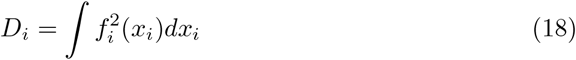

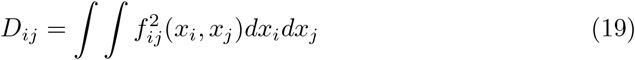

The sensitivity indices are:

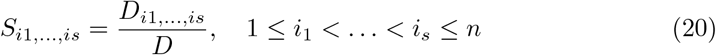

Thus, all terms add up to 1:

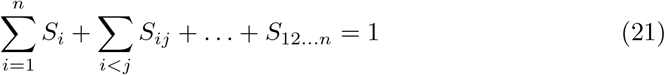

The primary impact of the input variable *x*_*i*_ on the output is determined by the first-order sensitivity index *S*_*i*_. In contrast, the impact of both *x*_*i*_ and *x*_*j*_ on the output is shown by the second-order sensitivity indices, and so forth. Each model result undergoes parameter ranking.

For this sensitivity analysis purpose, the Routh-Hurwitz stability criteria were adapted. Note that the following *λ* is the eigenvalues of the matrix **A**, where *d***x***/dt* = **Ax**.

#### Theorem 1

*A necessary and sufficient condition that the equation*

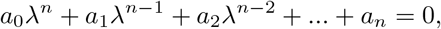

*with real coefficients in which the coefficient a*_0_ *is assumed to be positive, has only roots with negative real parts, is that the values of the following determinants all be positive [54, 55]*.

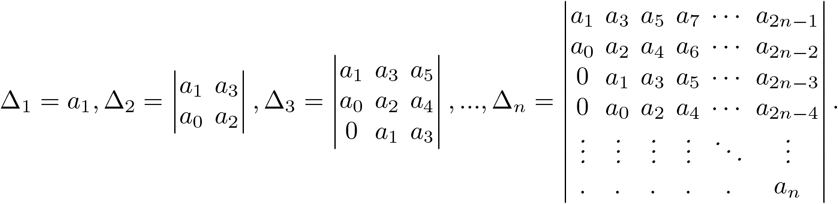

The characteristic equations for this study are as follows.

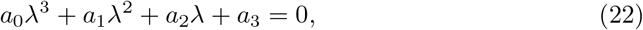

where

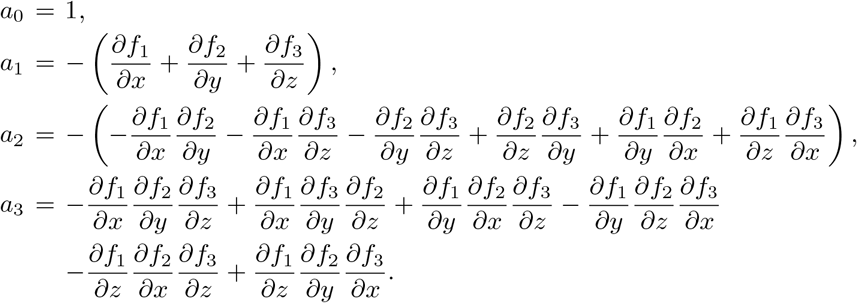

The system will be stable if the values of Δ are positive, where 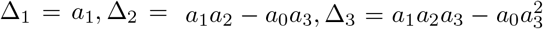. We adapted this criterion to obtain the sensitivity index.

## 3 Results

### 3.1 Smoker and non-smoker system states do not statistically differ

The demographics of the study cohort were presented in [26] (see Figure 1 in [26]). The average age of the sample population was 30 years, consisting of 90% males. For ethnicity, most were from the Middle East and North Africa (75%) followed by Asia (24%) and Africa (1%), with an average body mass index (BMI) of 25.9 Kgm^*−*2^. There were no significant differences between smokers and non-smokers for BMI. Smokers had a higher percentage of family members with cancer, hypertension, diabetes, and asthma.

As stated in Section 2.2, we processed our dataset with CCA and obtained a set of covariates as well as a reduced dimensionality system, where **X, Y**, and **Z** represent coarse states of psychological function, metabolism, and microbiome composition, respectively (as explained in Section 2.2). The descriptive statistics of the scaled covariates resulting from the data preprocessing step are shown in Table 1.

**Table 1:** The descriptive statistics of the covariates and Kruskal-Wallis mean rank comparison test between smokers and non-smokers

| Category | Variable | Mean | St. Dev. | Q1 | Q2 | Q3 | Min | Max |
| --- | --- | --- | --- | --- | --- | --- | --- | --- |
| Smoker | X | 0.5466 | 0.1948 | 0.3855 | 0.5322 | 0.6876 | 0.1771 | 1.0000 |
|  | Y | 0.6110 | 0.2185 | 0.4849 | 0.6312 | 0.7612 | 0.0000 | 1.0000 |
|  | Z | 0.5855 | 0.2248 | 0.4449 | 0.6155 | 0.7546 | 0.0000 | 0.9964 |
| Non-smoker | X | 0.5073 | 0.2031 | 0.3935 | 0.4612 | 0.6388 | 0.0000 | 0.9512 |
|  | Y | 0.5799 | 0.2074 | 0.4198 | 0.5769 | 0.6991 | 0.1546 | 0.9775 |
|  | Z | 0.5522 | 0.1816 | 0.4530 | 0.5524 | 0.6472 | 0.0543 | 1.0000 |

*Kruskal-Wallis mean rank comparison test (smokers vs. non-smokers): all p-values < 5%.*
*Note: X, Y, Z denote psychological traits, metabolism, and microbiome, respectively.*

According to Table 1, the mean value of the psychological traits covariates for smokers looks greater than that of non-smokers. On the other hand, the metabolic pathways and bacteria composition covariates for smokers seem to be less than those of non-smokers. However, the results of the Kruskal-Wallis mean rank test was not significant at the 5% level.

### 3.2 Feature Importance Analysis (FIA)

To identify the important features of each dataset, we employed radar plots based on the loadings (weights) of CCA, Figure 2 shows the top 10 most important features of psychological traits, metabolic pathways, and bacteria composition to maximize the correlation between variables. We also divided the data set into three criteria (all data, smokers, and non-smokers).

**Fig. 2:**
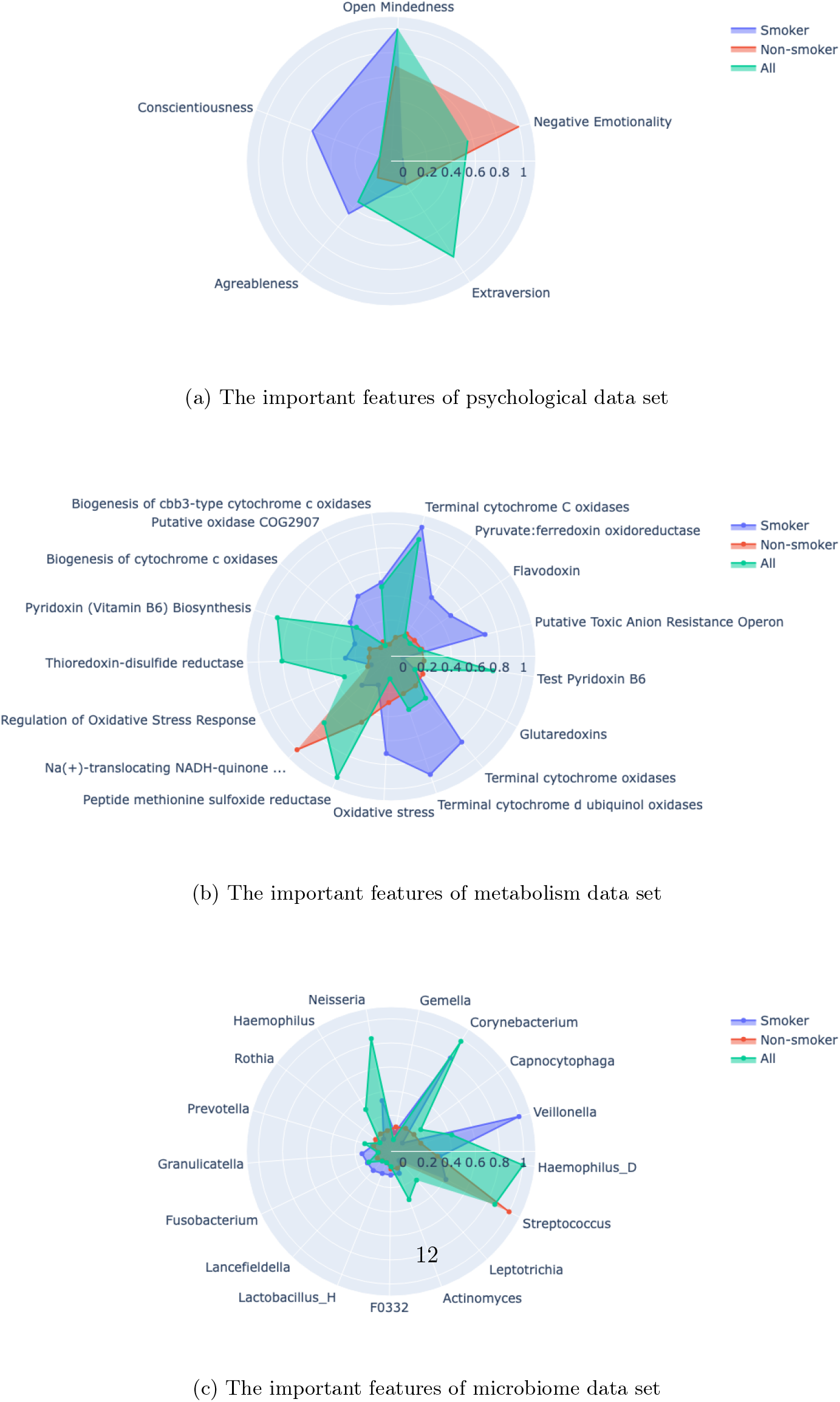
The important features of each variable

Open-mindedness, extraversion and negative emotionality are the most important features of the psychological set. *Peptide methionine sulfoxide reductase, Pyridoxin (Vitamin B6) Biosynthesis*, and *Terminal cytochrome C oxidases* are the three most important features for metabolism. In addition, *Haemophilus D, Corynebacterium*, and *Streptococcus* were the most essential features in the Bacteria set. Furthermore, the important characteristics of smokers and non-smokers were identified. Openness was identified as the primary feature in smokers, whereas negative emotionality was the most prominent feature for non-smokers. The difference is also noted in metabolic pathways. Three kinds of *Terminal cytochrome oxidases* (*Terminal cytochrome C oxidases, Terminal cytochrome d obiquinol oxidases, Terminal cytochrome oxidases*) followed by *Oxidative stress* are the most dominant features in smokers. Meanwhile, *Na(+)-translocating NADH-quinone oxidoreductase & rnf-like group of electron transport complexes* followed by *Peptide methionine sulfoxide reductase* and *oxidative stress* are the top three most important features for non-smokers. For microbiota, *Veillonella* and *Corynebacterium* genera are the top 2 primary features for smokers, while *Streptococcus* and *Haemophilus* genera are dominant in non-smokers.

In addition, to identify the difference between smokers and non-smokers, a summary of statistics of the ten most important features is presented in Table 2. Based on this table, the mean and median values of metabolism and microbiome were quite different, whereas the values were similar for the psychological data set. A Kruskal-Wallis test examined the mean rank difference between the two groups. The results showed that the mean rank of the psychological data set was not significantly different between smokers and non-smokers. Two features in the metabolism and microbiome data set resulted in significant differences between smokers and non-smokers. These are *Putative oxidase COG2907, Pyruvate:ferredoxin oxidoreductase* for metabolism; *Fusobacterium* and *Veillonella* genera for microbiome. Interestingly, the variables with the highest loading did not always confer significant statistical differences.

**Table 2:** Descriptive Statistics of the most important features for smokers and non-smokers

| Feature | Mean | Smoker<br>Med | Std | Mean | Non-Smoker<br>Med | Std | KW. | P-val |
| --- | --- | --- | --- | --- | --- | --- | --- | --- |
| **A. Psychological Traits** |  |  |  |  |  |  |  |  |
| Extraversion | 9.87 | 10 | 2.56 | 9.66 | 10 | 2.22 | 0.43 | 0.51 |
| Agreeableness | 11.53 | 12 | 2.14 | 11.54 | 12 | 2.46 | 0.09 | 0.77 |
| Conscientiousness | 11.40 | 11 | 3.57 | 10.92 | 11 | 2.70 | 0.12 | 0.73 |
| Negative Emotionality | 8.29 | 8 | 2.84 | 8.14 | 8 | 3.32 | 0.17 | 0.68 |
| Open Mindedness | 10.16 | 10 | 1.98 | 10.68 | 11 | 2.24 | 1.91 | 0.17 |
| **B. Metabolism** |  |  |  |  |  |  |  |  |
| Biogenesis of cbb3-type... | 25.25 | 8.00 | 54.68 | 20.76 | 10.33 | 27.53 | 1.14 | 0.29 |
| Biogenesis of cytochrome c... | 26.40 | 11.50 | 39.44 | 20.71 | 13.33 | 23.04 | 0.01 | 0.94 |
| Flavodoxin | 92.85 | 64.00 | 94.98 | 66.06 | 47.00 | 62.14 | 1.62 | 0.20 |
| Glutaredoxins | 339.15 | 204.38 | 315.73 | 264.16 | 187.95 | 247.87 | 0.91 | 0.34 |
| Na(+)-translocating NADH... | 285.28 | 160.00 | 353.11 | 219.98 | 124.00 | 258.34 | 0.24 | 0.62 |
| Oxidative stress | 267.51 | 175.04 | 258.28 | 195.84 | 138.67 | 177.38 | 1.28 | 0.26 |
| Peptide methionine sulfoxide... | 307.01 | 197.55 | 273.55 | 256.94 | 163.78 | 260.68 | 0.61 | 0.43 |
| Putative Toxic Anion... | 69.13 | 35.00 | 82.58 | 52.60 | 35.50 | 68.05 | 0.61 | 0.43 |
| Putative oxidase COG2907 | 12.60 | 5.00 | 36.17 | 6.44 | 2.17 | 11.65 | 4.90 | 0.03* |
| Pyridoxin (Vitamin B6)... | 195.64 | 115.62 | 182.42 | 147.29 | 106.78 | 134.62 | 1.39 | 0.24 |
| Pyruvate:ferredoxin... | 35.95 | 18.00 | 57.10 | 19.86 | 7.75 | 29.70 | 8.36 | 0.00* |
| Regulation of Oxidative Stress... | 29.85 | 14.67 | 40.98 | 23.46 | 11.00 | 36.56 | 0.18 | 0.67 |
| Terminal cytochrome C oxidases | 52.68 | 20.17 | 88.96 | 39.26 | 25.42 | 46.87 | 0.00 | 1.00 |
| Terminal cytochrome d... | 184.49 | 121.58 | 183.25 | 135.27 | 89.08 | 129.54 | 1.50 | 0.22 |
| Terminal cytochrome oxidases | 182.79 | 121.58 | 178.48 | 135.28 | 88.08 | 128.72 | 1.49 | 0.22 |
| Test Pyridoxin B6 | 183.00 | 114.91 | 175.72 | 131.40 | 93.60 | 117.12 | 1.87 | 0.17 |
| Thioredoxin-disulfide reductase | 162.88 | 98.25 | 160.48 | 120.74 | 83.07 | 108.65 | 1.26 | 0.26 |
| **C. Microbiome genus** |  |  |  |  |  |  |  |  |
| Actinomyces | 10855.98 | 2216 | 32476.60 | 6872.60 | 1699.0 | 13205.56 | 0.50 | 0.48 |
| Capnocytophaga | 3521.33 | 991 | 7012.86 | 4479.48 | 933.5 | 8814.91 | 0.00 | 0.95 |
| Corynebacterium | 9581.93 | 507 | 49565.89 | 5966.86 | 630.5 | 14719.40 | 0.04 | 0.84 |
| F0332 | 1330.98 | 30 | 6303.57 | 1395.28 | 26.5 | 5405.24 | 0.26 | 0.61 |
| Fusobacterium | 3737.22 | 1337 | 7425.76 | 1742.36 | 676.0 | 2457.02 | 3.88 | 0.04* |
| Gemella | 11012.62 | 3352 | 20934.93 | 9120.00 | 2506.0 | 16666.87 | 0.20 | 0.66 |
| Granulicatella | 4846.51 | 1867 | 7692.44 | 1840.54 | 1096.0 | 2084.31 | 2.28 | 0.13 |
| Haemophilus | 8108.76 | 2070 | 14138.17 | 5578.62 | 1763.0 | 10442.77 | 0.03 | 0.86 |
| Haemophilus_D | 21125.29 | 5568 | 33505.84 | 17399.34 | 3965.5 | 39498.24 | 0.28 | 0.59 |
| Lactobacillus_H | 782.09 | 2 | 4913.93 | 22.10 | 1.0 | 86.43 | 1.27 | 0.26 |
| Lancefieldella | 900.02 | 56 | 5513.99 | 187.28 | 42.5 | 727.88 | 3.76 | 0.05 |
| Leptotrichia | 2141.15 | 349 | 6720.78 | 1506.08 | 158.0 | 4453.55 | 2.72 | 0.10 |
| Neisseria | 13369.25 | 3113 | 32043.76 | 9120.82 | 4183.5 | 13908.52 | 1.76 | 0.19 |
| Prevotella | 5744.18 | 2670 | 9636.69 | 3511.96 | 1547.0 | 5209.23 | 2.95 | 0.09 |
| Rothia | 10269.58 | 4136 | 12774.52 | 6020.20 | 3421.0 | 6845.52 | 1.68 | 0.19 |
| Streptococcus | 133878.02 | 83113 | 125551.24 | 114904.00 | 69261.5 | 131348.01 | 0.77 | 0.38 |
| Veillonella | 16957.51 | 6165 | 42271.53 | 8587.66 | 2530.5 | 15400.31 | 3.98 | 0.04* |

\*significant at 5% level

Finally, we have further explored if the most important variable has the potential to discriminate smokers and non smokers. To do that, we employed several classification algorithms that allow for feature importance (See Appendix B for details). We concluded that smoking does not exhibit a strong signature within the most important features of our dataset since the accuracy is low (≈ 50%). This implies that differences in the above features do not allow us to distinguish smokers from non-smokers.

### 3.3 Metabolism and microbiome are highly interconnected and correlated

Here, we aimed to identify how the most important features interact. Figure 3 depicts the linear association network among the most important features of the variables and their degree of association (i.e. the number of connections made). These features were based on the most important features in Section 3.2 consisting of five features of Psychological traits, ten features of Metabolism, and ten features of Microbiome variables.

**Fig. 3:**
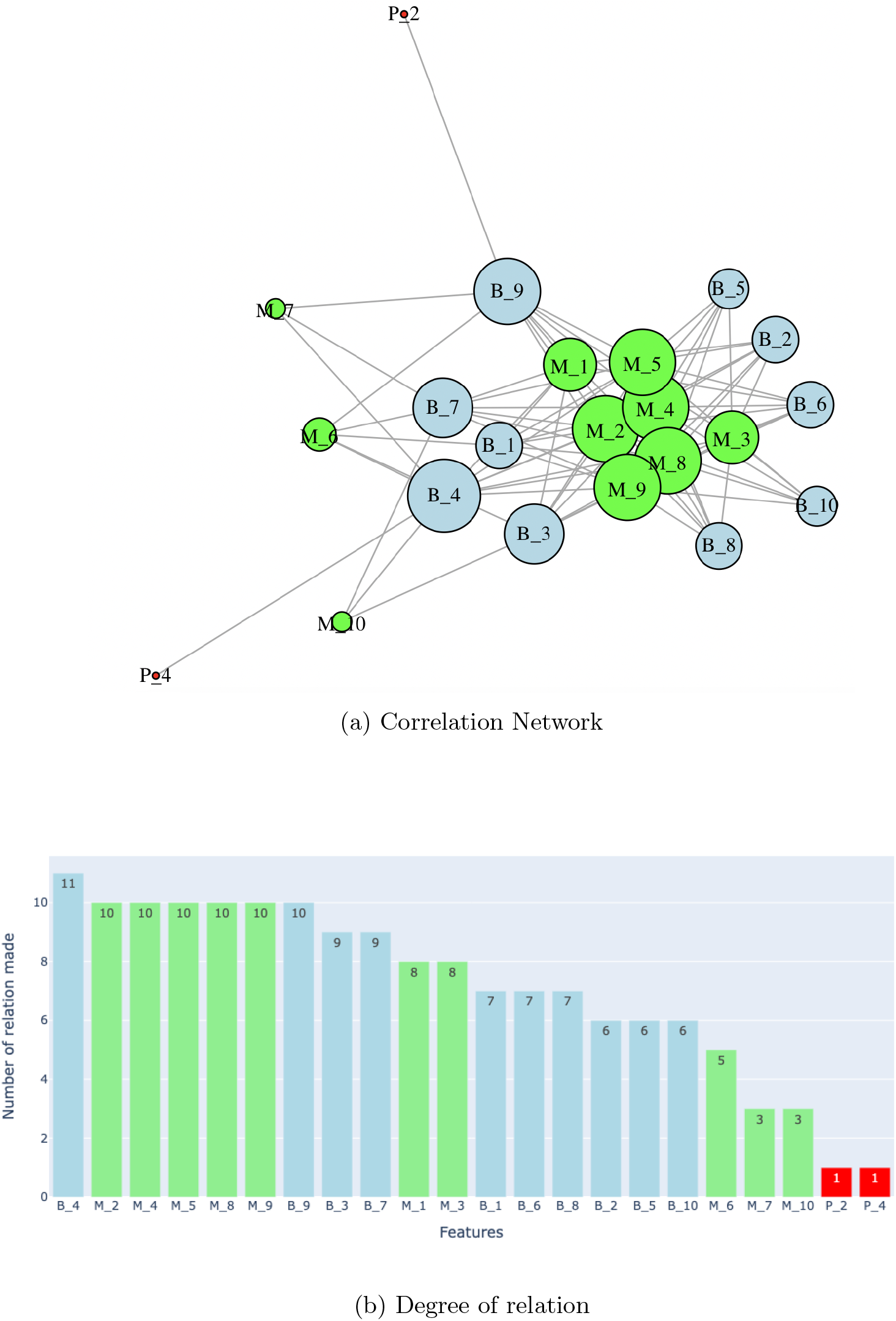
Correlation Network and the degree of relation for each feature

We denoted the sorted Psychological Traits features as *P*_1_, …, *P*_5_, then *M*_1_, …, *M*_10_ and *B*_1_, …, *B*_10_ for Metabolic Pathways and Bacteria Composition, respectively (see Table 3).

**Table 3:**
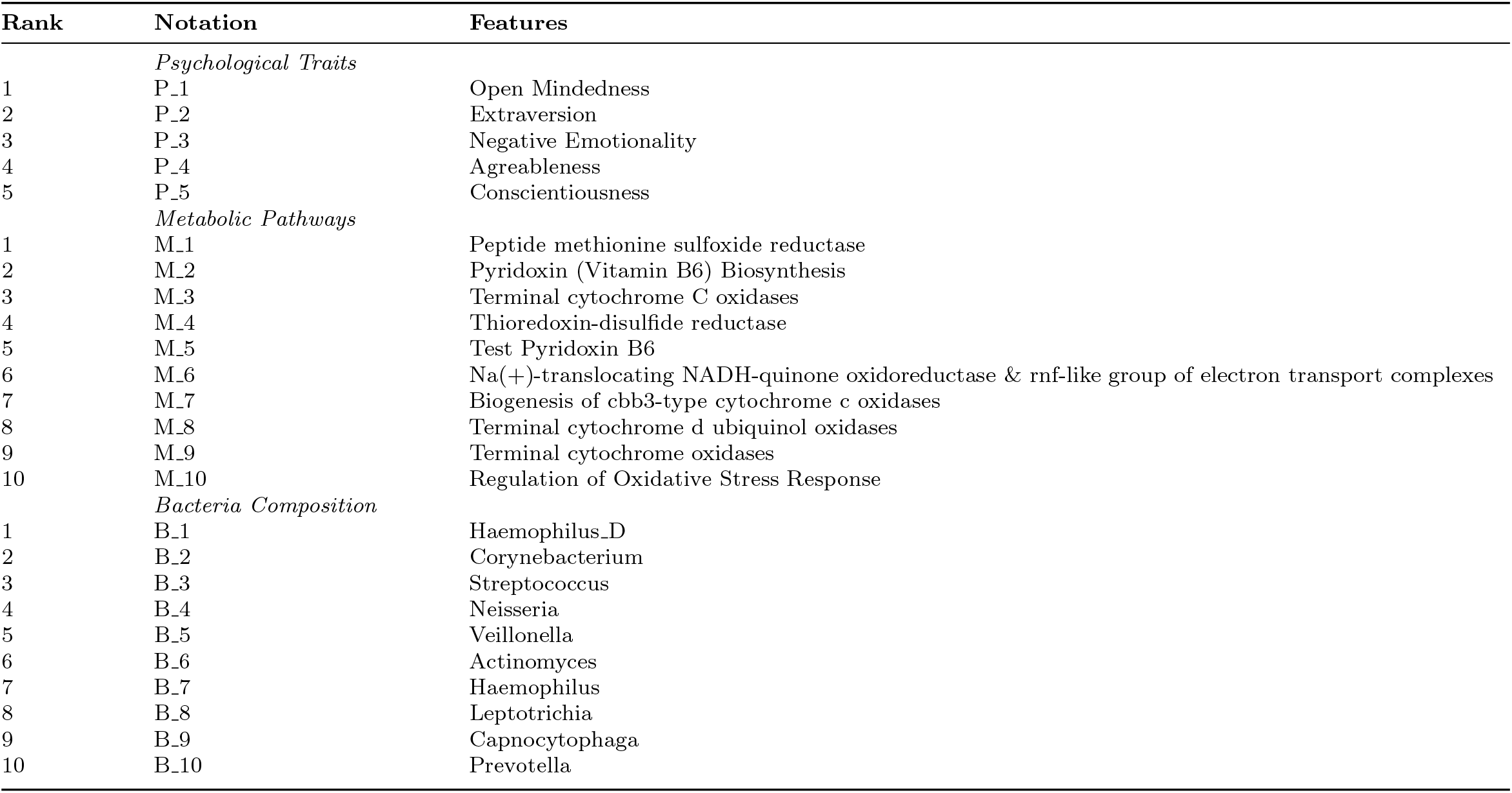
The details of the 10 important features of each variable

The largest node is the *Neisseria* genus (*B*_4_), which has a significant relationship with all metabolic features and the Agreableness feature. *Neisseria* species are identified as the fourth most widespread group of bacteria of adult oral microorganisms. Nonpathogenic *Neisseria* strains are considered important in maintaining oral and nasal environments by preventing potential pathogens from colonizing [56].

Moreover, the second largest nodes come from metabolic pathways and the Bacteria genus. Pyridoxin (Vitamin B6) Biosynthesis (*M*_2_), Thioredoxin-disulfide reductase (*M*_4_), Test Pyridoxin B6 (*M*_5_), Terminal cytochrome d ubiquinol oxidases (*M*_8_), and Terminal cytochrome oxidases (*M*_9_). These are significantly related to all Bacteria Composition features. On the other hand, the Capnocytophaga genus (*B*_9_) has a significant relation with all Metabolic features (except Regulation of Oxidative Stress Response) and the Extraversion feature. In general, metabolic pathways and bacterial composition are closely (linearly) related, while psychology seems to have a significant relationship with a few bacterial genera.

To enrich the knowledge of the relation among features in those three variables, a clustered correlation heat map was produced for all features (not only the ten most important features as above). Figure C1 (in the Appendix) shows three clusters of correlation (low, moderate, and high) among them. The details of the members of the cluster are given in Table C4 (see the Appendix). Interestingly, all psychological features were in the low cluster, suggesting they have a weak linear relation to metabolism and microbiome features.

### 3.4 Identification of steady state interactions for the coarse Brain-Metabolism-Microbiome system in smokers and non-smokers

In this section, our goal is to identify the interactions between the coarse system variables (CCA variates) at the steady state. To achieve this, we employed a regularized non-linear regression as described in Section 2.4. The results are provided in Figure 4 (see Table D5 in the Appedix for further details).

**Fig. 4:**
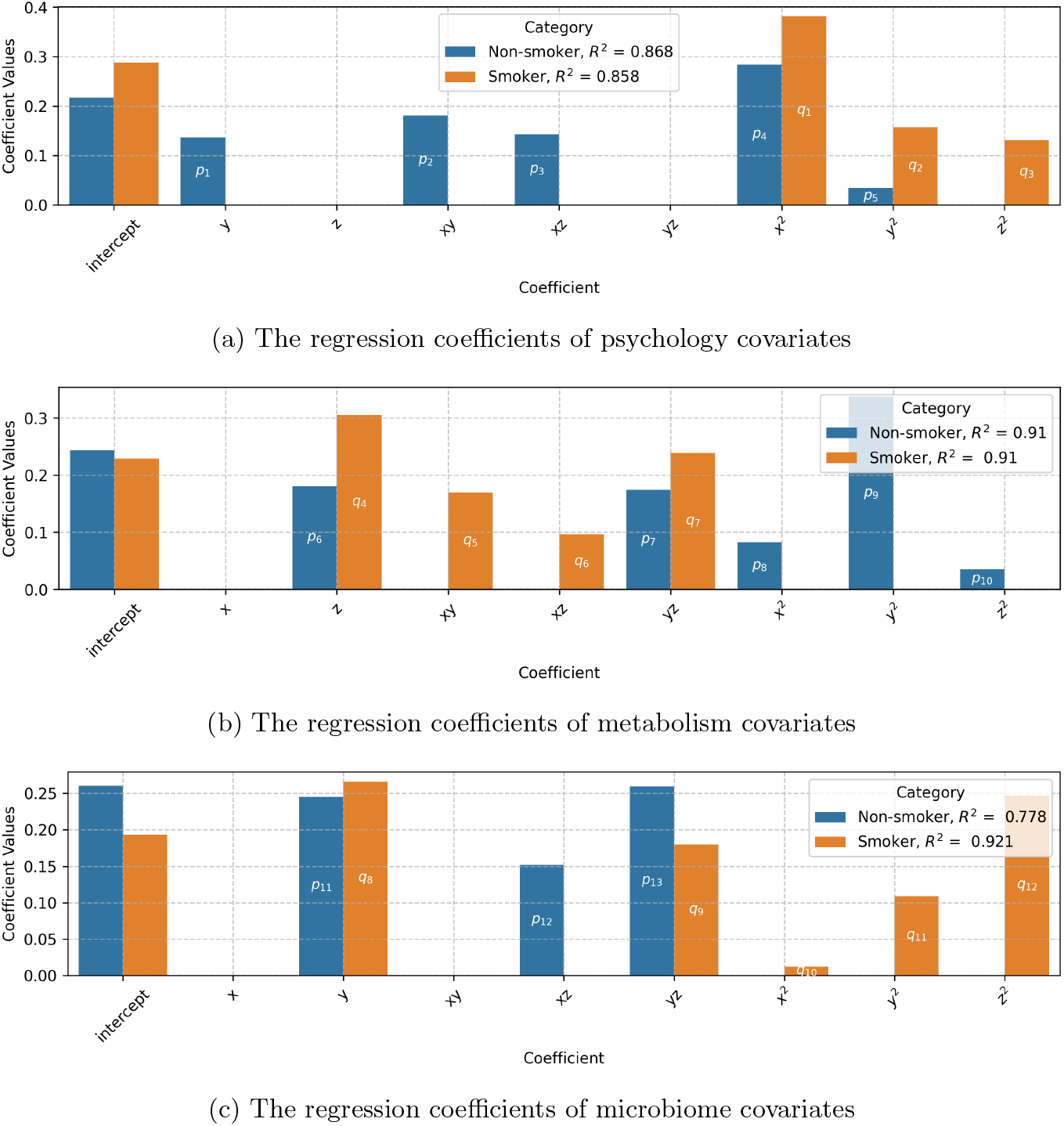
The regression coefficients of smokers and non-smokers category along with their coefficient of determination (*R*^2^) for each variable. *p* and *q* are the notation of parameters for sensitivity analysis in Section 3.4.

The regression coefficients in Table D5 show the causal relationship between the variables seen from the regression coefficients. This is further explained in Figure 5, which illustrates the relation among the three variables differentiated by smoking status. Metabolites and bacteria are tightly connected, and we represent them as a block, which in turn interacts with the brain state.

**Fig. 5:**
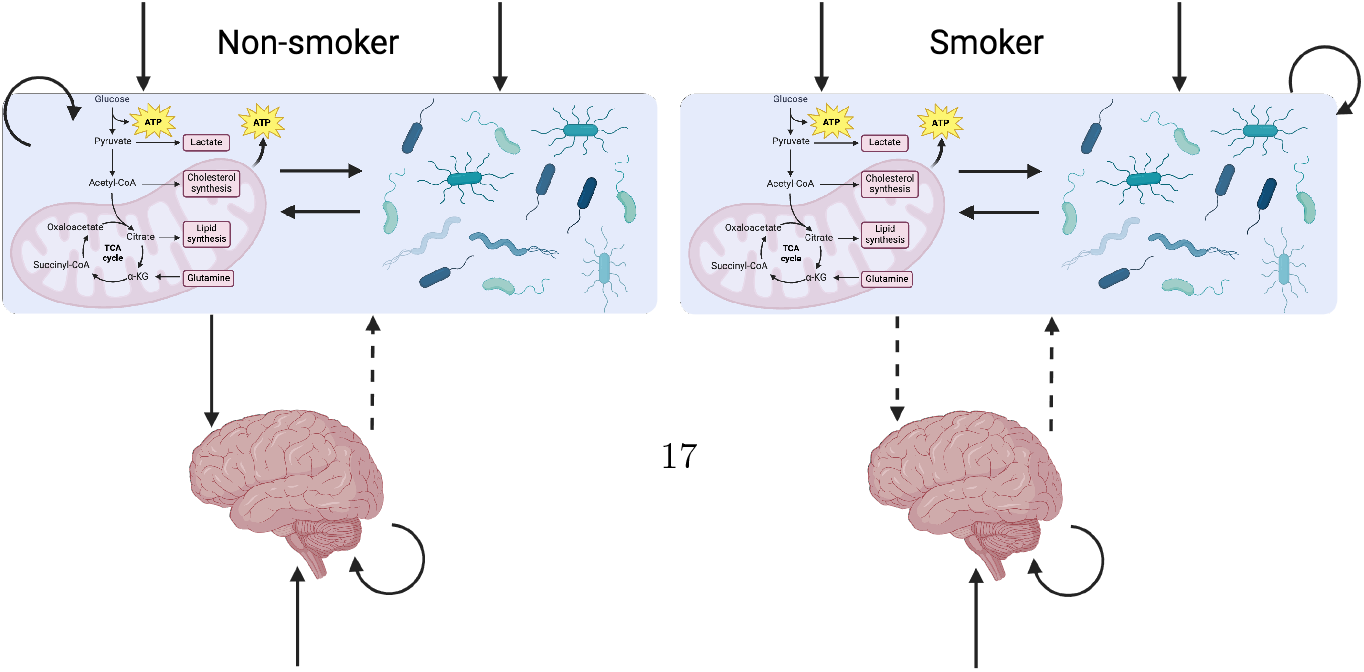
Identified interaction diagram between Brain, Metabolism and Microbiota (Created with BioRender.com). Solid lines denote linear interactions and dashed ones the non-linear interactions.

Based on Figure 5, one can see that the big difference is in terms of direct effects. Smoking can weaken the relationship between the brain and the Metabolism-Microbiome (MM) block, by eliminating the direct effect only from the brain to the MM block. A potential reason for the weak connection between brain and MM is due to the timescale separation, where the psychological (behavioral) aspect changes in quite longer period than metabolism and microbiome.

The analysis identified a positive feedback network between the three systems (represented by their respective covariates), which is perturbed by smoking.

#### Robustness of the healthy system state and potential routes to pathology

Here, we aimed to understand the properties of the steady state interaction regarding its robustness. The idea is that healthy (non-smoking) steady state should be stable and robust upon parameter perturbation. Pathologies should emerge upon changing the interaction coefficient, combinations and destabilizing the steady states. The latter can provide hints for medical interventions.

The stable equilibrium points obtained using parameters in Table D5 (in the Appendix) are *x*_*s*_ = 0.4019, *y*_*s*_ = 0.4417, and *z*_*s*_ = 0.4073 for smokers, and *x*_*ns*_ = 0.3804, *y*_*ns*_ = 0.4436, *z*_*ns*_ = 0.4467 for non-smokers. These stable point values are within one standard deviation of the mean value of the scaled data for non-smokers. Thus, no statistical difference exists for the (*x, y, z*) state between non-smokers and smokers. This validates our assumption that the mean value of a non-smoker should be a stable fixed point for a significant range of perturbations. In the following, we study the robustness of the steady state using sensitivity analysis. Based on the stable points, smokers have slightly higher values of the psychological variables.

#### Sensitivity analysis and stability properties

We are now interested in investigating how this stable point can be unstable, particularly what interaction, when perturbed, leads to associated pathologies. Recalling the regression coefficients in Figure 4, the parameters of the non-smoker equations are denoted by *p*_1_, …, *p*_13_ while *q*_1_, …, *q*_12_ are for smokers (see Table E6 in the Appendix).

In this analysis, the parameters were perturbed to ±80% of their values. Besides, we set the steady-states as the initial values. The results of the interaction coefficients sensitivity analysis are visualized in Figure E2 (see the Appendix). In general, for non-smokers, the three main parameters that are the most sensitive are the self-feedback loop of metabolism (*y*^2^), the self-feedback loop of psychological term (*x*^2^), and the interaction between metabolism and microbiome (*yz*) in affecting microbiome. The system becomes unstable when these parameters are perturbed by 385%, 458%, and 1888% of their original values, respectively. Those three variables are nonlinear terms in the equation. This means that nonlinear effects are the most sensitive parameters to destabilize the system of non-smokers.

Moreover, the most sensitive parameters for smokers are the self-feedback loop of psychological variable (*x*^2^), the self-feedback loop of microbiome (*z*^2^), and the interaction between metabolism and microbiome in influencing metabolism (*yz*). These coefficients will destabilize the system if their values are perturbed more than 366%, 1213%, and 965% of the original values, respectively. This result aligns with the non-smokers case where the nonlinear terms play a significant role in destabilizing the steady state. Figure 6 highlights the three most sensitive connections for smokers and non-smokers. In turn, we assume that the least perturbation parameters are more probable to destabilize the system. As a result the most probable destabilization factor for non-smokers is the self-feedback loop of metabolism, whereas for smokers, it is the self-feedback loop of psychological traits. Regarding interactions, the metabolism-microbiome interaction is the most sensitive parameter, but its direction differs between smokers and non-smokers. In non-smokers, this interaction primarily affects the microbiome, whereas in smokers, it primarily affects metabolism.

**Fig. 6:**
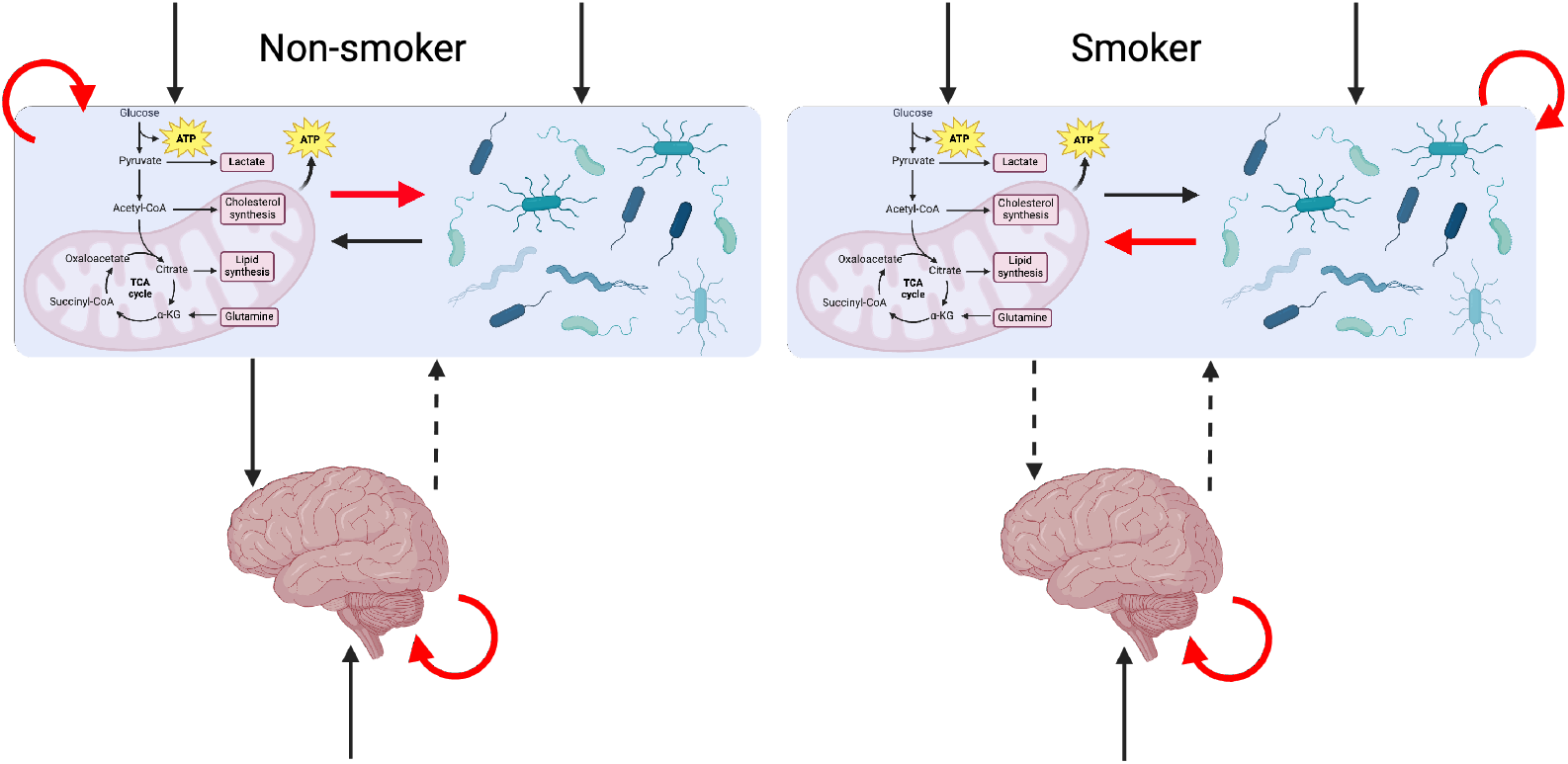
Red interactions indicate the ones that are able to destabilize the observed steady states of the BMM system (Created with BioRender.com). Solid lines denote linear interactions and dashed ones the non-linear interactions.

## 4 Discussion

In this study, we sought to investigate how the oral microbiota, metabolic pathways, and psychological characteristics among smokers are connected. A statistical analysis comparing the mean values of variable covariates failed to capture the differences between smokers and non-smokers. This is because these factors do not act as isolated variables linked to smoking but rather exert a collective effect as a system. Consequently, these interactions are not fully reflected in traditional statistical tests.

A correlation network of the most important features across the three variables provides a visual mapping of their interactions. The *Veillonella* genus was identified as a feature with a moderate degree of relation and the only significant factor differentiating smokers from non-smokers. According to [57], *Veillonella* was one of the genera found abundantly in healthy smokers. This bacterium also showed an increased presence in the oral microbiome of smokers [58]. To better manage smokers with particular behavioral patterns, the identified microbial biomarkers could lead to new and more effective clinical applications. Further research is warranted to investigate the oral microbiome’s composition and its functional relationships with mental health and cigarette smoking.

The central finding of our work is the interaction network between the oral microbiome, metabolic activity, and psychological traits in the context of the oral-brain axis. In particular, we discovered that there is (i) a mutual activation between the microbiome and metabolism and (ii) a positive feedback cascade between these subsystems. This result is in accordance with the Selfish Brain theory, where such positive feedback interactions are assumed between metabolism and the brain [59, 60]. Of course, this is not the complete picture. Rainer Straub has extended the concept by postulating that there is a competing behavior between the immune system and the brain for energy resources [61, 62, 63]. Therefore, if we had the corresponding immune profiles of our samples, we would have a more comprehensive picture of the system dynamics.

Our results indicate that smoking interferes with the interacting dynamics between the oral microbiome, metabolic activity, and psychological traits. In particular, smoking plays a role in the relationship between psychology and metabolism. The linear effect from the oral microbiome to the brain can be weakened for smokers. In this case, smoking may potentially interfere with psychiatric treatment (e.g., depression) [64, 65]. Thus, a personalized treatment for smokers is needed. Note that the direction of the effect needs further examination in future work. Furthermore, when the interaction between covariates was analyzed within a system dynamics framework, a distinction between smokers and non-smokers became evident. However, this difference was not observed when comparing their mean values. (see Section 3.1).

This work has some limitations, including the size and origin of the dataset, which only covers the UAE, which limits the generality of the results. Limited sample size has implications in the generality of important features as derived by the CCA loadings, which can be unstable for small samples [66]. Moreover, our FIA is related to the specific target of the CCA method i.e. correlation and dimensionality reduction. Future studies may consider additional data and further analytics such as the partial least square (PLS) method. In addition, our FIA results for psychological traits may be further explored to define psychological types (profiles) related to smoking in future studies.

Using the nonlinear regression model, we can capture only partially the BMM system. Firstly, we only consider the nonlinear regression *up to the second order*, and the corresponding noise term represents the potential structural uncertainties. On the other hand, other organs influence the BMM system, as for instance the immune system, which are not captured in the current study (as discussed above).

Our data only represent a very sparse time resolution (single time point) of the investigated dynamics. To overcome this limitation, we assume that the expected values of our data represent the steady-state of the BMM variables, although it is generally known that living systems are poised out of equilibrium [67]. Therefore, we need state that our findings represent only *potential* dynamics of the system. Longitudinal data would help to identify system dynamics more comprehensively and with sufficient confidence.

Finally, we expect the existence of a *multiscale* behavior of the system, which is not fully considered in the current study. In particular, there is a discernible timescale separation between the psychological changes and microbiome-metabolism dynamics. Being internal to the body, the latter can exhibit rapid changes, while psychological changes can unfold over weeks or months [68, 69]. Therefore, Singular Perturbation Analysis may be considered in future work to provide further insights.

## Declarations

### Ethics approval and consent to participate

Not applicable.

### Consent for publication

Not applicable.

### Availability of data and materials

The datasets used and/or analysed during the current study are available from Dr. Mohammad T. Albataineh on reasonable request.

### Authors’ contributions

SMU and HH developed the methodology. SMU implemented the methodology and wrote the paper under the supervision of HH, HJ, SS, and MTA. SS assisted with sensitivity analysis. MTA provided data and with HH defined the research questions. All authors reviewed and provided feedback.

## Acknowledgments

HH have received funding from the BMBF under grant agreement No. 031L0237C (MiEDGE project/ERACOSYSMED). HH would also like to acknowledge the support of the Volkswagen Stiftung for the “Life?” initiative (grant number 96732). Finally, HH and SS acknowledges the support of the RIG-2023-051 grant from Khalifa University and the UAE-NIH Collaborative Research grant AJF-NIH-25-KU. We extend our thanks to Prof. Anna-Maria Pappa for her help. This work utilized ChatGPT to enhance the quality of English.

## Appendix A The Detailed Features in Section 2

Table A1 describes Metabolic Pathways while Table A2 depicts the genera of Bacteria Composition.

## Appendix B Smoking Status Prediction

Smoking status was predicted using features from BMM variables with a Random Forest classifier optimized by XGBoost. The accuracy was evaluated using 3-fold cross-validation (see Table B3).

## Appendix C The Clustered Correlation Heat Map

Figure C1 and Table C4 present the clusters of the correlation between the features.

**Table A1:**
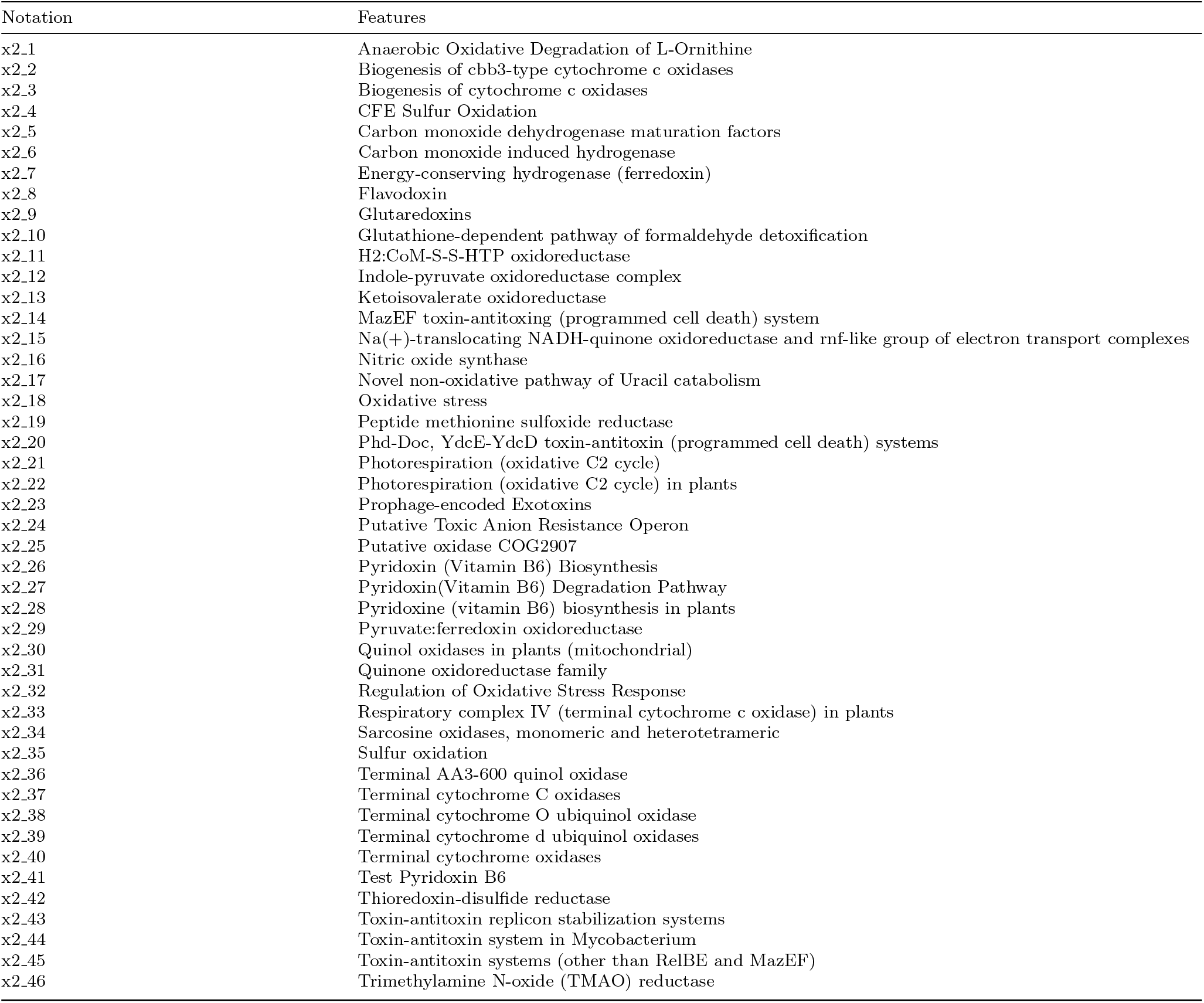
The details of Metabolic Pathways Features

**Table A2:**
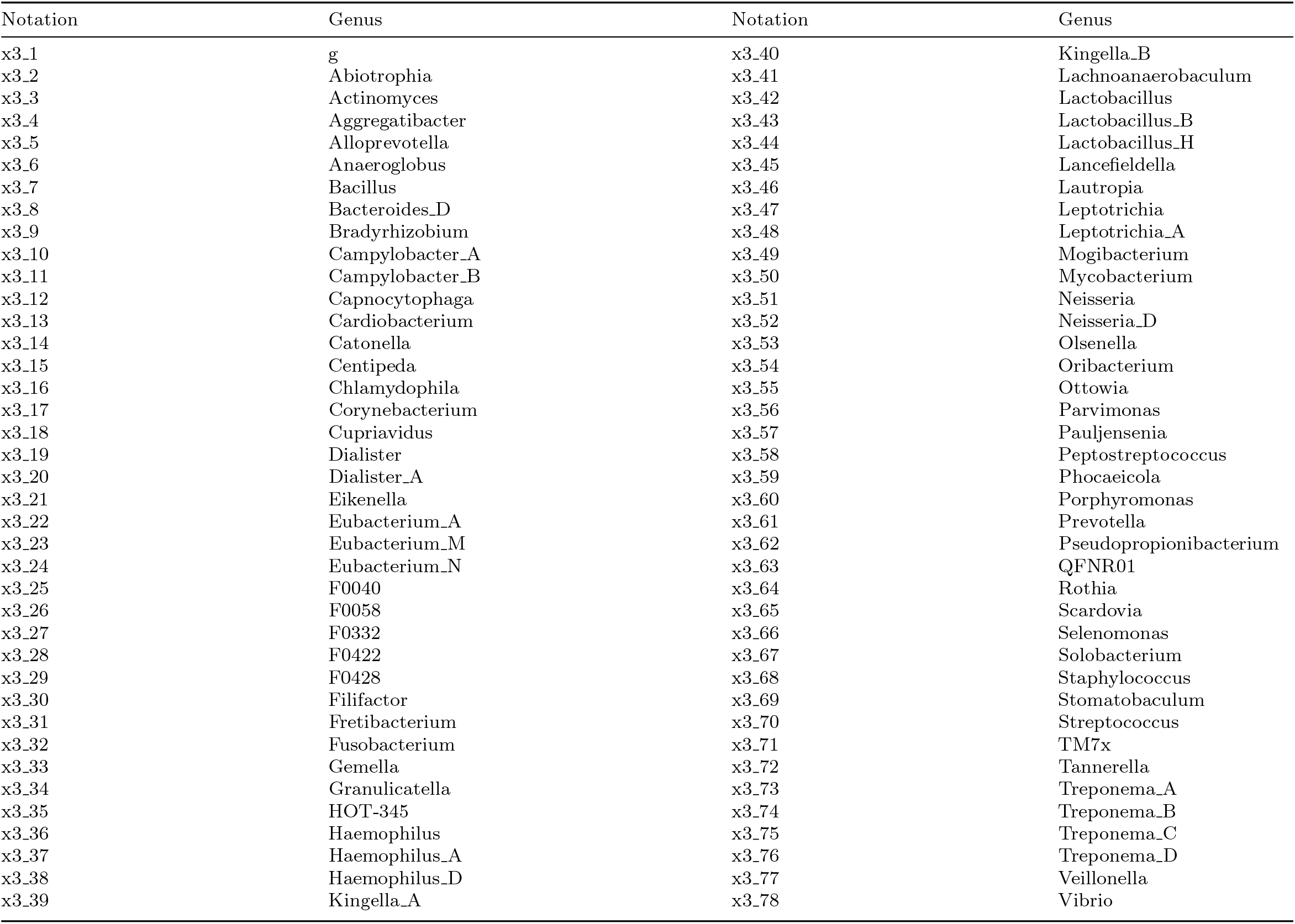
The genera in Bacteria Composition

**Table B3:**
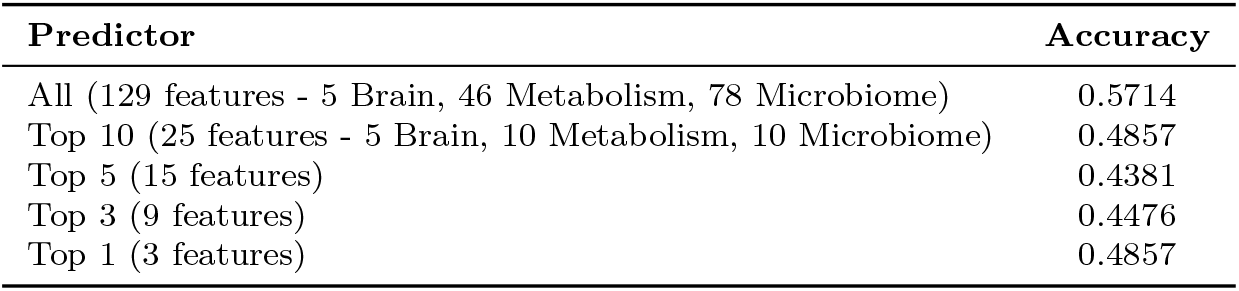
Smoking status classification results

**Fig. C1:**
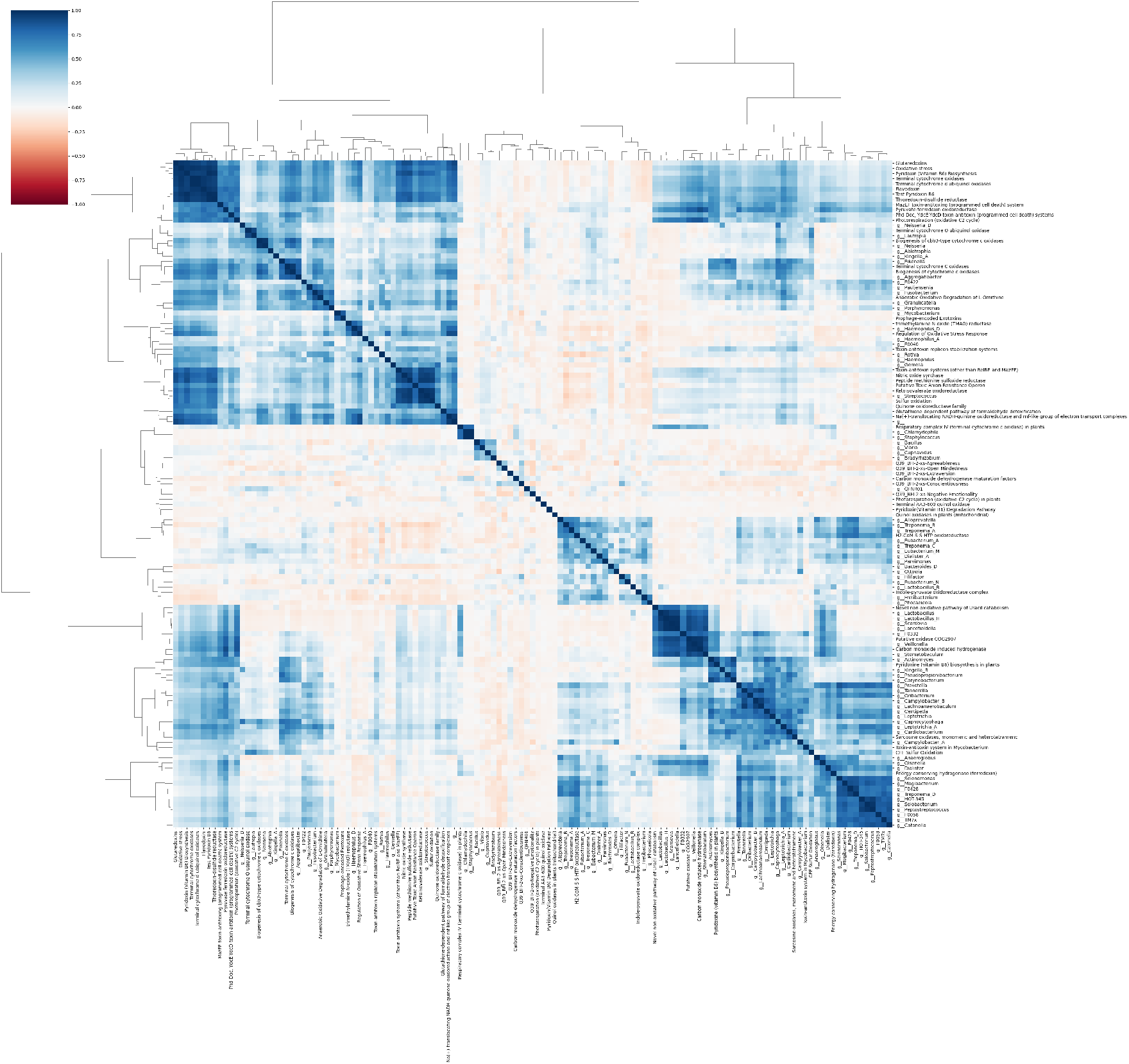
The clustered correlation heat map for all features

**Table C4:**
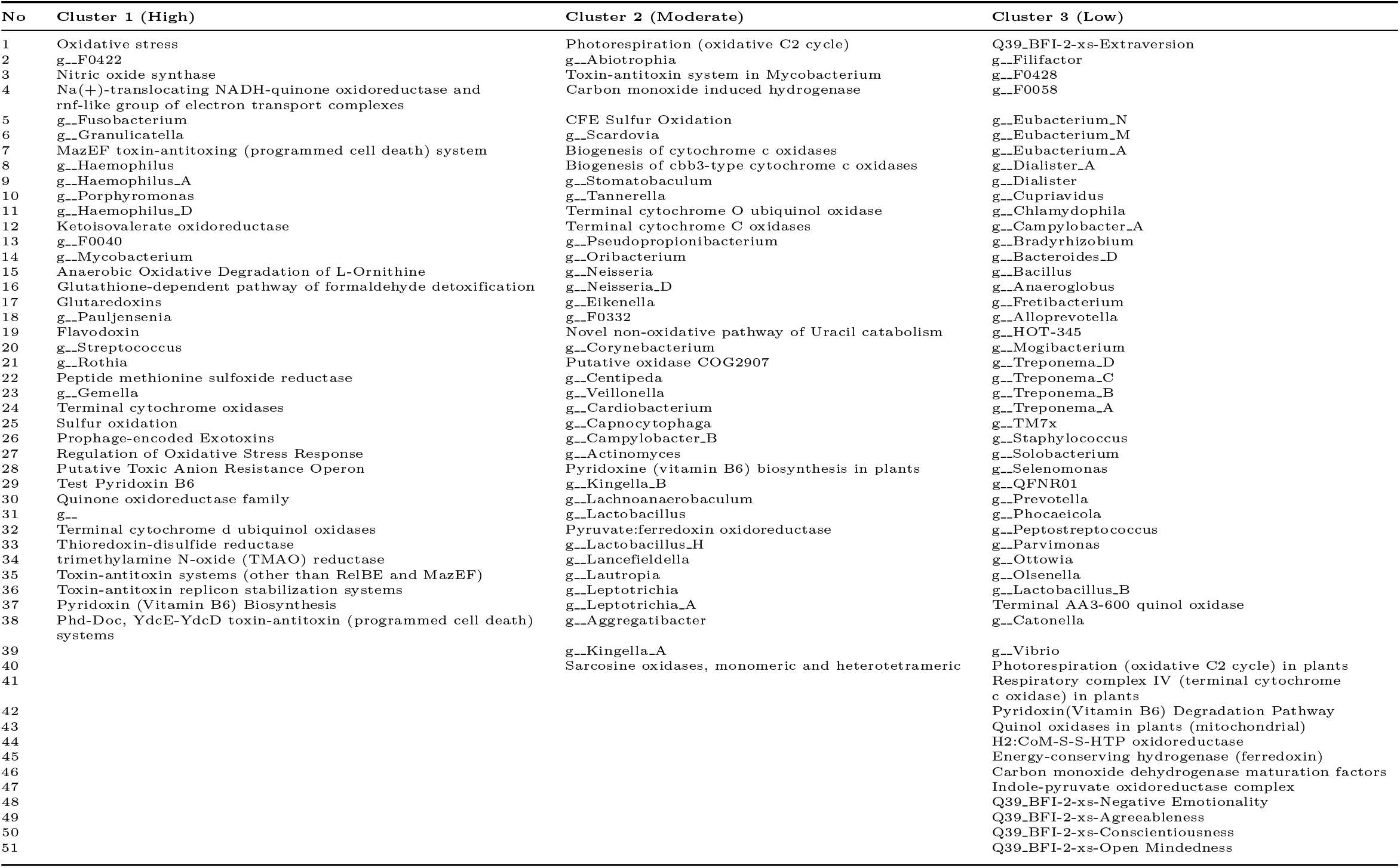
The details of the clustered correlation of the heat map

## Appendix D Nonlinear Regression

The details of the parameter estimates of nonlinear regression for smokers and non-smokers is given in Table D5.

**Table D5:** The parameter estimates of nonlinear sparse regression

| $i$ | $b_{i,0}$ | $b_{i,1}$ | $b_{i,2}$ | $b_{i,3}$ | $b_{i,4}$ | $b_{i,5}$ | $b_{i,6}$ | $b_{i,7}$ | $b_{i,8}$ | $R^2$ |
| --- | --- | --- | --- | --- | --- | --- | --- | --- | --- | --- |
| *Non-smoker* |  |  |  |  |  |  |  |  |  |  |
| x | 0.2173 | 0.1365 | 0 | 0.1807 | 0.1426 | 0 | 0.2838 | 0.0342 | 0 | 0.868 |
| y | 0.2433 | 0 | 0.1806 | 0 | 0 | 0.1738 | 0.0823 | 0.3369 | 0.0352 | 0.91 |
| z | 0.2606 | 0 | 0.2454 | 0 | 0.1518 | 0.2595 | 0 | 0 | 0 | 0.778 |
| *Smoker* |  |  |  |  |  |  |  |  |  |  |
| x | 0.2879 | 0 | 0 | 0 | 0 | 0 | 0.382 | 0.1571 | 0.1312 | 0.858 |
| y | 0.2287 | 0 | 0.3052 | 0.1697 | 0.096 | 0.2381 | 0 | 0 | 0 | 0.91 |
| z | 0.1932 | 0 | 0.2663 | 0 | 0 | 0.1795 | 0.0118 | 0.109 | 0.2473 | 0.921 |

## Appendix E Sensitivity analysis

The parameter notation for sensitivity analysis is presented in Table E6 while the sensitivity indexes for each parameter of smokers and non-smokers is depicted in Figure E6.

**Table E6:** The parameter notation for sensitivity analysis

| $i$ | $b_{i,0}$ | $b_{i,1}$ | $b_{i,2}$ | $b_{i,3}$ | $b_{i,4}$ | $b_{i,5}$ | $b_{i,6}$ | $b_{i,7}$ | $b_{i,8}$ |
| --- | --- | --- | --- | --- | --- | --- | --- | --- | --- |
| *Non-smoker* |  |  |  |  |  |  |  |  |  |
| x | 0.2173 | $p_1$ | 0 | $p_2$ | $p_3$ | 0 | $p_4$ | $p_5$ | 0 |
| y | 0.2433 | 0 | $p_6$ | 0 | 0 | $p_7$ | $p_8$ | $p_9$ | $p_{10}$ |
| z | 0.2606 | 0 | $p_{11}$ | 0 | $p_{12}$ | $p_{13}$ | 0 | 0 | 0 |
| *Smoker* |  |  |  |  |  |  |  |  |  |
| x | 0.2879 | 0 | 0 | 0 | 0 | 0 | $q_1$ | $q_2$ | $q_3$ |
| y | 0.2287 | 0 | $q_4$ | $q_5$ | $q_6$ | $q_7$ | 0 | 0 | 0 |
| z | 0.1932 | 0 | $q_8$ | 0 | 0 | $q_9$ | $q_{10}$ | $q_{11}$ | $q_{12}$ |

**Fig. E2:**
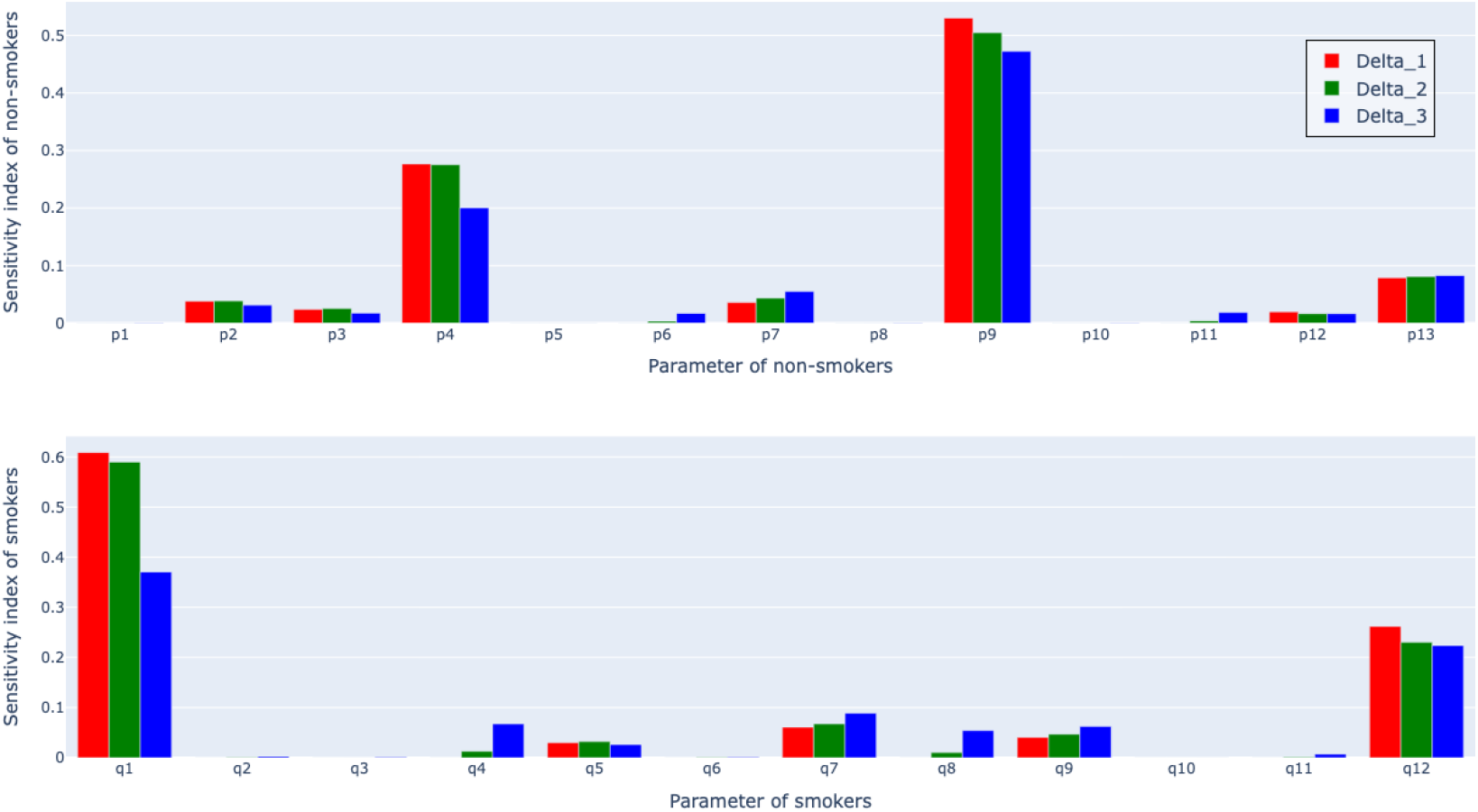
The sensitivity index of the parameter for non-smokers and smokers

